# Enhancing HIV-1 neutralization by increasing the local concentration of MPER-directed bnAbs

**DOI:** 10.1101/2022.10.23.513422

**Authors:** Soohyun Kim, Maria V. Filsinger Interrante, Peter S. Kim

## Abstract

Broadly neutralizing antibodies (bnAbs) against the membrane-proximal external region (MPER) of the gp41 component of the HIV-1 envelope (Env) are characterized by long hydrophobic heavy-chain complementarity-determining regions (HCDR3s) that interact with MPER and some viral membrane lipids, to achieve increased local concentrations. Here, we show that increasing the local concentration of MPER-directed bnAbs at the cell surface via binding to the high affinity Fc receptor (FcγRI) potentiates their ability to prevent viral entry in a manner analogous to the previously reported observation whereby the lipid binding activity of MPER bnAbs increases their concentration at the viral surface membrane. However, binding of MPER-directed bnAb 10E8 to FcγRI abolishes the neutralization synergy that is seen with the N-heptad repeat (NHR)-targeting antibody D5_AR and NHR-targeting small molecule enfuvirtide (T20), possibly due to decreased accessibility of the NHR in the FcγRI-10E8-MPER complex. Taken together, our results suggest that lipid-binding activity and FcγRI-mediated potentiation function in concert to improve the potency of MPER-directed bnAbs by increasing their local concentration near the site of viral fusion. Therefore, lipid-binding may not be a strict requirement for potent neutralization by MPER-targeting bnAbs, as alternative methods can achieve similar increase in local concentration while avoiding potential liabilities associated with immunologic host tolerance.

**Author summary:** The trimeric glycoprotein Env is the only viral protein expressed on the surface of HIV-1, is the target of broadly neutralizing antibodies, and is the focus of most vaccine development efforts. Broadly neutralizing antibodies targeting the membrane proximal external region (MPER) of Env show lipid-binding characteristics and modulating this interaction affects neutralization. In this study, we tested the neutralization potencies of variants of the MPER-targeting antibody 10E8 with different viral membrane-binding and host FcγRI-binding capabilities. Our results suggest that binding to both lipid and FcγRI improves the neutralization potency of MPER-directed antibodies by concentrating the antibody at sites of viral fusion. As such, lipid-binding may not be uniquely required for MPER-targeting broadly neutralizing antibodies, as alternative methods to increase local concentration can achieve similar improvements in potency.

## Introduction

Despite 40 years of extensive research, human immunodeficiency virus (HIV) remains a major global public health concern, with more than 1.5 million new cases in 2020 and 38 million people currently living with HIV/acquired immunodeficiency syndrome (AIDS) (https://www.who.int/news-room/fact-sheets/detail/hiv-aids). HIV-1 infection is initiated by binding of the viral envelope glycoprotein (Env), a trimer consisting of the gp120 and gp41 subunits, to CD4 on CD4^+^ T cells facilitated by cellular coreceptors (e.g., CXCR4 or CCR5), to trigger membrane fusion (1, 2). During fusion, Env undergoes a substantial conformational change, forming a prehairpin intermediate (PHI) (3–6), in which the N-heptad repeat (NHR), C-heptad repeat (CHR), and membrane-proximal external region (MPER) of gp41 are exposed, all of which are poorly accessible on the native pre-fusion Env protein (7–10).

Broadly neutralizing antibodies (bnAbs) targeting the MPER have remarkable breadth, neutralizing > 98% of primary HIV-1 isolates (11, 12). The long hydrophobic heavy-chain complementarity-determining region 3s (HCDR3s) of MPER-directed antibodies bind lipid components of the viral membrane through electrostatic interactions with anionic phospholipids, which is reported to enhance the activity of anti-MPER bnAbs (2F5 and 4E10) in blocking viral infection (9, 13–17). Crystal structures of 4E10 in complex with lipids phosphatidic acid (PA), phosphatidylglycerol (PG) and glycerol phosphate revealed that the HCDR1 and HCDR3 loops of 4E10 interact with polar lipid head and hydrophobic lipid tail groups, respectively (18). Likewise, the light chain of 10E8, another anti-MPER bnAb, was predicted by X-ray crystallography and cryo-electron microscopy to bind the Env-membrane interface (8, 19, 20), while crystal structures of 10E8 complexed with its epitope scaffold and either PG or PA identified the 10E8 lipid-binding sites within the LCDR1 and HCDR3 loops (19). Hence, lipid-binding is an important consideration for the development of successful MPER-based immunogens. However, effective induction of bnAbs by vaccine immunogens is limited due to the autoreactivity to host lipids conferred by the conserved viral epitopes (21). Initially, 10E8 was reported to show no autoreactivity towards lipids (22), but later found to bind weakly to membranes, with preference for those that are cholesterol-rich (15, 19, 23, 24). For 10E8, decreasing and increasing its electrostatic interaction with the viral membrane diminished and enhanced its neutralization potency, respectively, while for 4E10, the correlation between interactions with lipids and neutralization potency was variable (15, 23, 24).

Another mechanism to modulate the neutralization potency of anti-MPER bnAbs is through FcγRI (CD64)-mediated potentiation. Previous work has shown that the neutralization potencies of anti-MPER bnAbs such as 2F5, 4E10, 10E8 and LN01 are enhanced as much as 5,000-fold in target cells expressing FcγRI (25–27). This potentiation has been attributed to the binding of the Fc portion of immunoglobulin (IgG) molecules to host FcγRI, allowing those antibodies to accumulate at the cell surface and increase the local concentration of antibodies able to block viral infection, potentially also providing a kinetic advantage that may be beneficial for bnAbs with epitopes only exposed for a short time during fusion (25).

Here, we evaluated the relationship between lipid-binding activity and FcγRI-mediated potentiation of anti-MPER bnAbs. We discovered that the two mechanisms modulate neutralization in a similar manner, both increasing local antibody concentrations at either the viral membrane or target cell membrane, leading to an increase in neutralization potency. We demonstrate that each mechanism can compensate for decreases in neutralization caused by loss of the other. Consistent with prior reports (28, 29), we show that increasing the local antibody concentrations of D5_AR, a monoclonal antibody that targets the NHR, increases neutralization potency while maintaining neutralization synergy with 10E8. On FcγRI-expressing cells, however, neutralization synergy was not symmetric for 10E8 and D5_AR, suggesting that the NHR epitope may have decreased accessibility in FcγRI-10E8-MPER complexes as compared to those formed in the absence of FcγRI. In summary, while lipid-binding activity of anti-MPER bnAbs improves neutralization by concentrating antibodies on the viral membrane, this effect can be accomplished by other means, suggesting that lipid-binding activity may not be a prerequisite for the high potency of MPER-targeting bnAbs.

## Results

### Generation of 10E8 variants with alterations in lipid-binding activity

To explore the relationship between lipid-binding activity of 10E8 and FcγRI-mediated potentiation, we produced 10E8 variants with both increased and decreased lipid-binding by modifying the interaction between antibody and lipid bilayers, as described previously (19, 23). As only the 10E8 heavy chain was reported to interact with MPER (19), to limit any effects on binding to MPER, we selected only variants with mutations within the light chain. We refer to variants with decreased and increased lipid-binding activity as 10E8d and 10E8i, respectively. 10E8d used in this paper is equivalent to “mutant 5” previously generated by Irimia et al (19), where two PG-or PA-binding residues within the LCDR1 were mutated to alanine (Fig. 1A,B). 10E8i is equivalent to “10E8-3R” previously generated by Rujas et al. (23), where three residues in the light chain that were predicted by X-ray crystallography and cryo-electron microscopy to contact the Env-membrane interface were substituted to positively charged arginines (Fig. 1A, B) to increase the electrostatic interaction with lipid bilayers, as measured by binding to vesicles containing 1,2-dioleoyl-sn-glycero-3-phosphocholine (DOPC) and 1,2-dioleoyl-sn-glycero-3-phosphoserine (DOPS) (8, 19, 20). To ensure that these variants did not affect binding to MPER, we generated 10E8 variants as monovalent fragment antigen-binding (Fab) fragments (Fig. S1) and tested their affinities against MPER. 10E8i Fabs showed similar (0.99-fold), while 10E8d Fabs showed slightly decreased (3.9-fold) affinity (K_D_) to MPER (Table 1), compared to 10E8 Fabs.

**Fig. 1.**
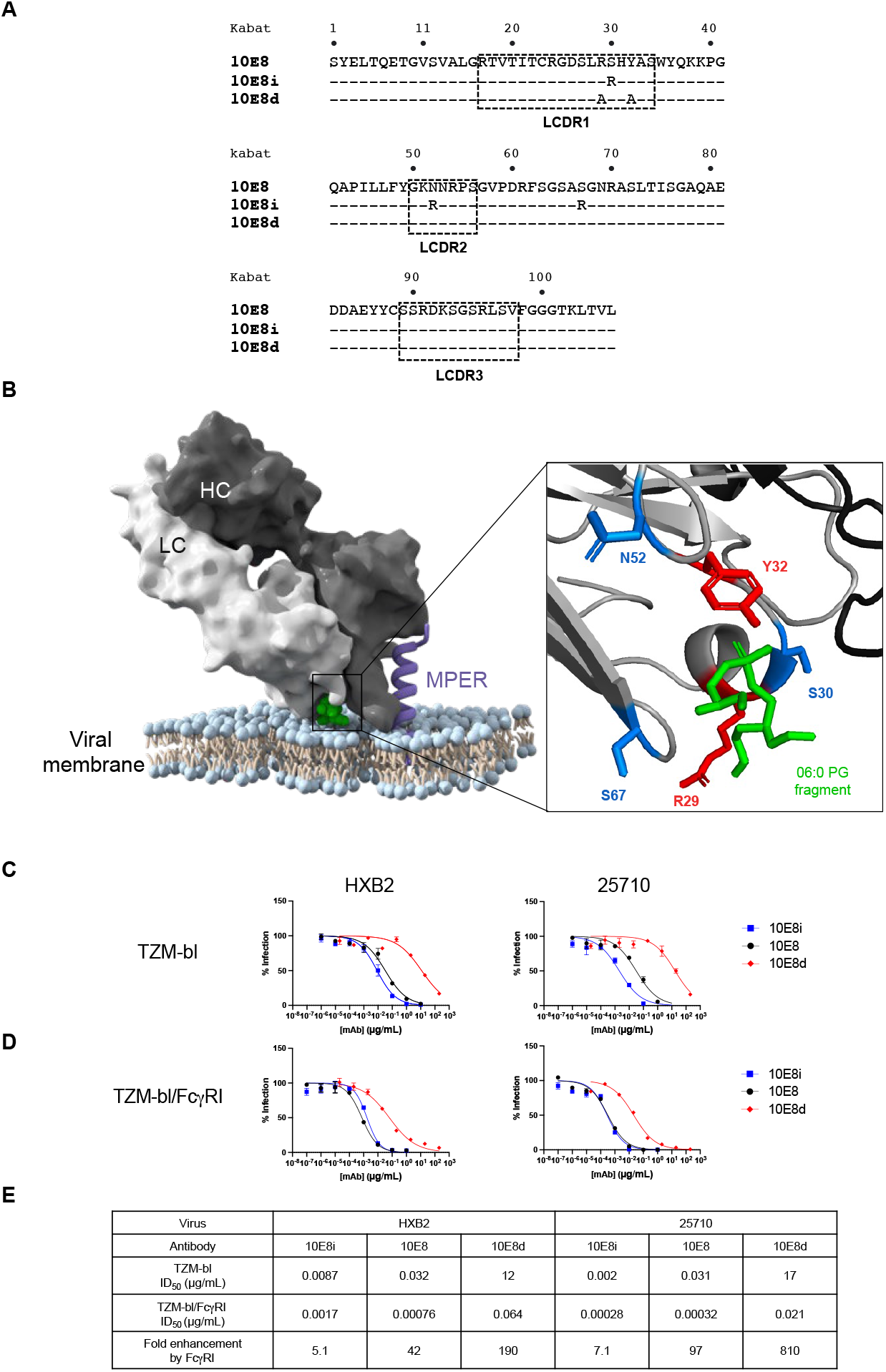
Altering local concentrations of 10E8 affects neutralization. (A) Alignment of light chain amino acid sequences of 10E8 variants are shown with Kabat numbering scheme on top of the alignment. Both 10E8i and 10E8d are variants identified and characterized elsewhere: 10E8d refers to mutant 5 from (19) and 10E8i refers to 10E8-3R from (23). (B) Model of 10E8 Fab in complex with the lipid and MPER (PDB: 5T85). The heavy chain (HC) and light chain (LC) of 10E8 are shown in dark gray and light gray, respectively. Position of MPER epitope (purple) and phosphatidylglycerol lipids (06:0 PG, green) are marked. (Right) Closeup view of residues and side chains altered in the variants on 10E8 to generate 10E8d (red) and 10E8i (blue) are marked. (C) Neutralization curves demonstrating changes in potency of 10E8 variants with increased (10E8i) or decreased (10E8d) lipid-binding activity against lentiviruses pseudotyped with HIV-1 HXB2 (tier-1B) and 25710 (tier 2C) in TZM-bl cells. Results are shown as mean ± standard error of the mean (SEM) from duplicates of two independent experiments. (D) Neutralization curves demonstrating FcγRI-mediated potentiation and lipid-binding activity-mediated changes in neutralization against lentiviruses pseudotyped with HIV-1 HXB2 and 25710 in TZM-bl cells/FcγRI. Results are shown as mean ± SEM from duplicates of two independent experiments. (E) Shown are the values of the antibody dose that reduces viral infection to 50% (ID_50_) and the fold-enhancement of neutralization potencies of 10E8 variants against lentiviruses pseudotyped with HIV-1 HXB2 and 25710 in TZM-bl cells versus TZM-bl/FcγRI cells. Fold enhancement is calculated by dividing the ID_50_ against TZM-bl cells by the ID_50_ against TZM/FcγRI cells.

**Table 1.**
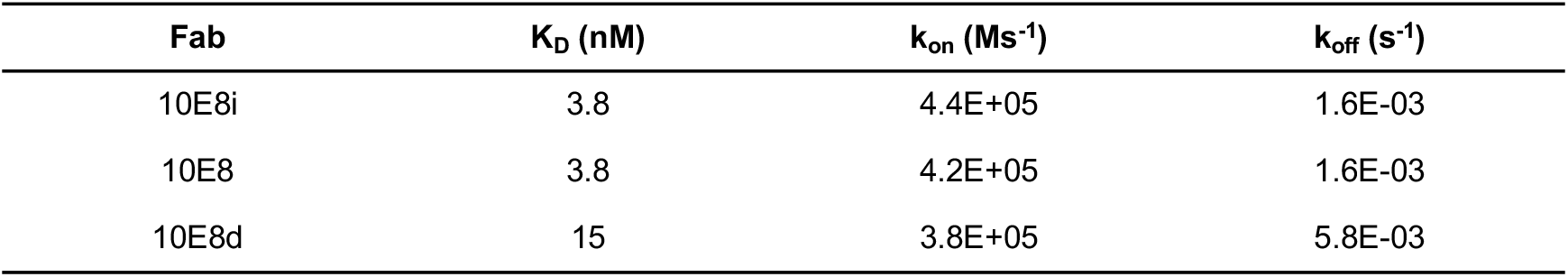
Affinity of 10E8 variants to biotinylated MPER peptide

### FcγRI-mediated potentiation of 10E8 variants is increased with decreased lipid-binding

To confirm that alterations in the lipid-binding activity of 10E8 affect neutralization, we tested the neutralization activity of 10E8 and its variants against HXB2 (tier 1B) and 25710 (tier 2C) HIV-1 pseudoviruses using Tat-regulated luciferase reporter gene expression in TZM-bl cells (30). As expected, the neutralization potencies of 10E8i and 10E8d increased and decreased, respectively, compared to 10E8 against both pseudoviruses based on the measured ID_50_ (Fig. 1C). Next, we tested the neutralization potencies of 10E8 and its variants against cells that express FcγRI. The increase in neutralization conferred by 10E8i was further increased in FcγRI-expressing cells (Fig. 1D). However, in the presence of FcγRI, the neutralization potency of 10E8 was similar to that of 10E8i, such that the enhanced neutralization potency conferred by FcγRI was limited (Fig. 1E). These results suggest that lipid-binding activity and FcγRI-mediated potentiation of 10E8 act similarly to concentrate antibodies at either the viral membrane or the target cell membrane to increase local antibody concentrations, and that there is a “ceiling” on the extent that the local concentration increases neutralization.

### 10E8 and D5_AR targeting the NHR of gp41 are synergistic

Some inhibitors that bind to gp41 that are more exposed in the PHI have been reported to show synergy when tested in combination (28, 31, 32). For instance, 10E8 was found to be synergistic with a mimetic of the gp41 CHR, C34, that targets the gp41 NHR and with the NHR mimic, 5-helix, which targets the CHR (28). Also, the MPER-targeting antibody 2F5 is synergistic with the NHR-targeting antibody D5 (29).

To investigate the effect of concentrating antibodies on neutralization synergy, we first tested candidate antibodies for their synergistic behavior, including 10E8v4, a variant of 10E8 that has been optimized for solubility (33), and D5_AR, a potency-improved variant of D5 (34). We used isobologram analyses of neutralization curves to evaluate the degree of synergy (35–37) of these two antibodies and found that the combination of 10E8v4 and D5_AR showed a synergistic effect on neutralization potency against HIV-1 pseudotypes tested in TZM-bl cells (Fig. 2A).

**Fig. 2.**
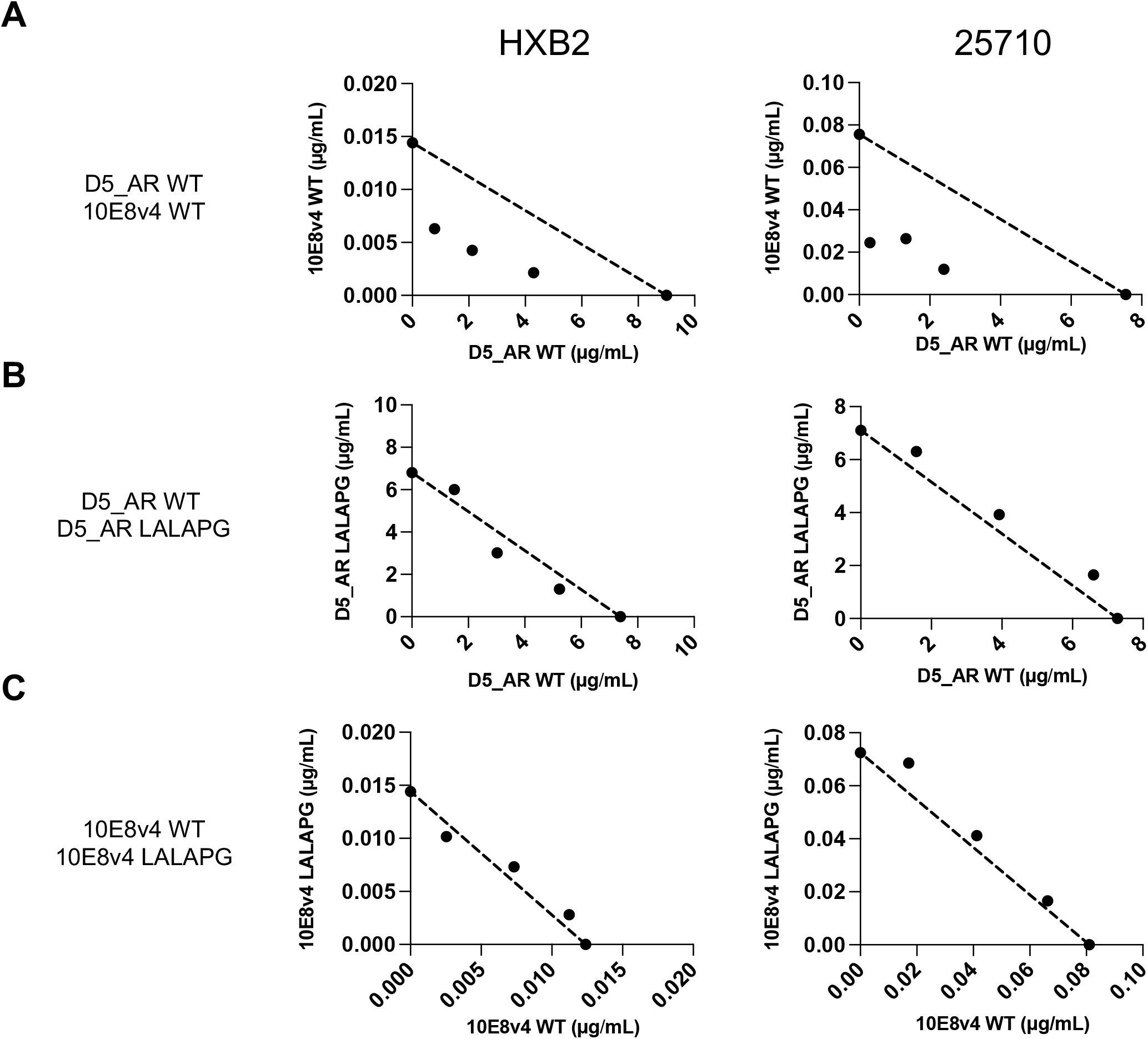
D5_AR and 10E8v4 are synergistic. Isobologram analyses of the combination of (A) D5_AR WT and 10E8v4 WT, (B) D5_AR WT and D5_AR LALAPG, and (C) 10E8v4 WT and 10E8v4 LALAPG against viruses pseudotyped with Env from HIV-1 strains HXB2 (left) and 25710 (right) in TZM-bl cells. The dotted lines indicate lines of additivity. Results are shown as mean ID_50_ performed in duplicates (technical replicates). Similar results were obtained in an independent repeat experiment. Data points below, along and above the line of additivity indicate synergy, additivity and antagonism, respectively.

### Effect of increasing local antibody concentrations on neutralization synergy

We used two strategies to modulate antibody concentrations at the site of viral fusion to assess the effect of concentrating antibodies on neutralization synergy. The first was FcγRI-mediated potentiation to increase the antibody concentrations at the cell surface. For the second, we used a LALAPG mutant that incorporates L234A, L235A, and P329G mutations within the Fc portion of the wild-type (WT) antibodies, known to block the Fc-Fc receptor interaction (38), to selectively modulate individual antibody concentrations in combination experiments. We first confirmed that the LALAPG mutations do indeed abolish binding of the D5_AR and 10E8v4 antibodies to FcγRI. While both D5_AR and 10E8v4 WT antibodies bound dose-dependently to recombinant FcγRI, both LALAPG mutants could not bind to FcγRI, as measured by enzyme-linked immunosorbent assay (ELISA) (Fig. S2A). Using flow cytometry, we confirmed that D5_AR and 10E8v4 WT and LALAPG mutants did not bind to the TZM-bl cells that were used in the neutralization assay. The WT antibodies bound to TZM-bl cells expressing FcγRI but the LALAPG mutants abolished this binding by D5_AR and 10E8v4 antibodies (Fig. S2B). We also confirmed that the LALAPG mutation abolishes FcγRI-mediated potentiation (Fig. S3 and Table S1). Since the WT and LALAPG mutants possess the same Fab, they should bind additively to their epitopes. Indeed, in control experiments where we tested combinations between D5_AR WT and D5_AR LALAPG, or 10E8v4 WT and 10E8v4 LALAPG, we saw additive effects in the isobologram analyses (Fig. 2B-C). These results suggest that the LALAPG mutation only affects binding between Fc and Fc receptor and that isobologram analyses are effective in assessing antibody synergy.

We next tested the combination of D5_AR LALAPG and 10E8v4 LALAPG in the presence of FcγRI as a control. Consistent with the synergy displayed between D5_AR WT and 10E8v4 WT in the absence of FcγRI (Fig. 2A), we found that combining D5_AR LALAPG and 10E8v4 LALAPG was also synergistic in FcγRI-expressing cells (Fig. 3A). We also observed synergy between D5_AR WT and the 10E8v4 LALAPG mutant (Fig. 3B). Similarly, we used 10E8v4 WT to increase local concentration of 10E8v4, but found quite unexpectedly that the synergy between D5_AR and 10E8v4 was abolished by the FcγRI-mediated increase in the local concentration of 10E8v4 (Fig. 3C). We also tested the combination of 10E8d, which has decreased neutralization potency compared to 10E8, and D5_AR. We also found that D5_AR and 10E8d were synergistic in the absence of FcγRI and the synergy was abolished in FcγRI-expressing cells (Fig. S4). These results suggest that formation of the FcγRI-10E8-MPER complex decreases the accessibility of the NHR in this case, to limit binding by D5_AR.

**Fig. 3.**
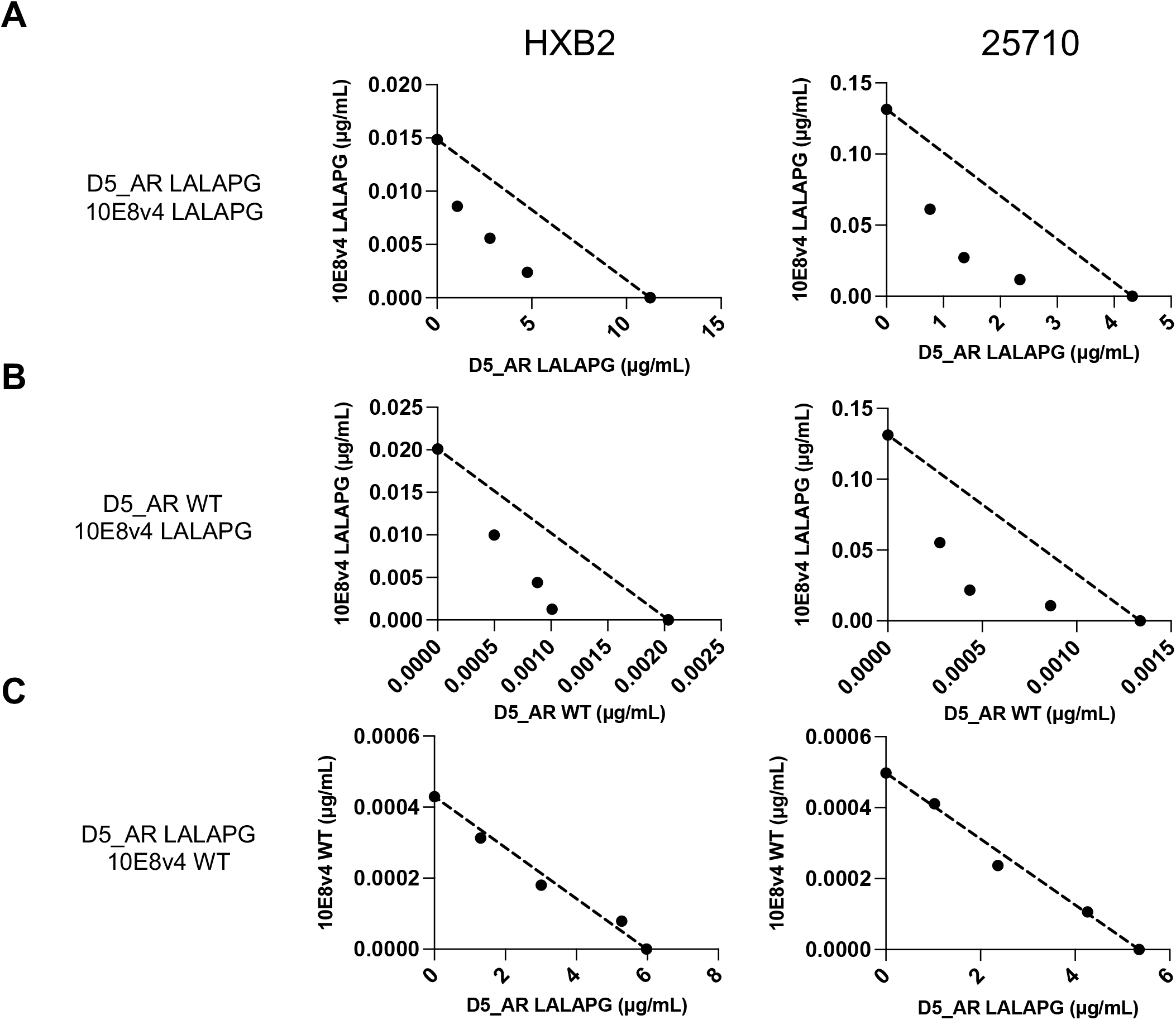
Relationship between neutralization synergy and FcγRI-binding. Isobologram analyses of the combination of (A) D5_AR LALAPG and 10E8v4 LALAPG, (B) D5_AR WT and 10E8v4 LALAPG and (C) D5_AR LALAPG and 10E8v4 WT against viruses pseudotyped with Env from HIV-1 strains HXB2 (left) and 25710 (right) in TZM-bl/FcγRI cells. The dotted lines indicate lines of additivity. Results are shown as mean ID_50_ performed in duplicates. Similar results were obtained in an independent repeat experiment. Data points below, along and above the line indicate synergy, additivity, and antagonism, respectively.

We then tested whether we could observe synergy with NHR-targeting agents other than D5_AR, which binds to the hydrophobic pocket of the NHR (39). We used enfuvirtide (T20), an FDA-approved peptide inhibitor of HIV-1 that binds outside the hydrophobic pocket of the NHR (40–42), and tested its combination with 10E8v4 in the presence and absence of FcγRI. Enfuvirtide was synergistic with 10E8v4 not bound to FcγRI (Fig. 4A-B), but this synergy was abolished when 10E8v4 was bound to FcγRI (Fig. 4C), analogous to the diminished synergy seen between 10E8 variants and D5_AR (Fig. 3C and Fig. S4).

**Fig. 4.**
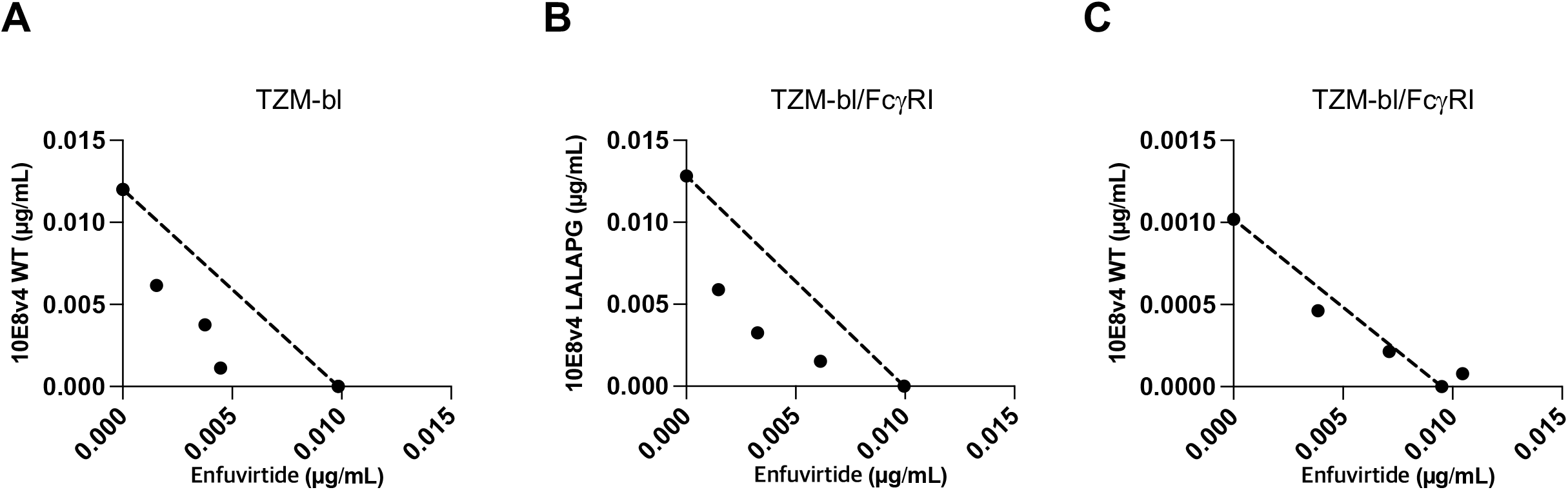
FcγRI-bound 10E8v4 abolishes synergy with an anti-NHR targeting agent, enfuvirtide. (A) Isobologram analyses of neutralization by the combination of enfuvirtide and 10E8v4 WT against viruses pseudotyped with Env from HIV-1 strain HXB2 in TZM-bl cells. Isobologram analyses of the combination of (B) enfuvirtide and 10E8v4 LALAPG and (C) enfuvirtide and 10E8v4 WT against viruses pseudotyped with Env from HIV-1 strains HXB2 in TZM-bl/FcγRI cells. The dotted lines indicate lines of additivity. Results are shown as mean ID_50_ performed in duplicates. Similar results were obtained in a separate repeat experiment, not shown here. Data points below, along and above the line of additivity indicate synergy, additive and antagonism, respectively.

## Discussion

Here, in agreement with earlier studies (13–17, 19, 23, 24), we show that increasing the local concentration of anti-MPER bnAbs correlates with enhanced neutralization potency (Fig. 1). However, there seems to be an upper limit or neutralization potency ceiling to the extent that the local concentration increases neutralization, as 10E8i (which was more potent than 10E8 in TZM-bl cells) and 10E8 were similarly potent against cells expressing FcγRI (Fig. 1). We hypothesize that the neutralization ‘potency ceiling’ we observe here might be due to a limit on the amount of antibody that can be physically present locally at the site of viral membrane fusion, due to FcγRI availability and/or steric effects.

Lipid-binding is important for enhancing neutralization of 10E8 and the specific roles we propose are depicted in Fig. 5. It was previously suggested that the lipid-binding properties of anti-MPER bnAbs increase their neutralization potencies through a two-step model where the antibodies first attach to the viral membrane, increasing the local antibody concentrations, followed by binding to Env (13, 15, 17, 19, 23, 43). In addition to this two-step model, the interaction between anti-MPER bnAbs and membrane has been proposed to enhance neutralization concomitant with or at a step after the engagement of Env, further stabilizing the antibody-Env complex (23). Our results show that even without increased lipid -binding activity, increasing local concentration via FcγRI can also separately improve neutralization potencies. Therefore, lipid-binding may not be a prerequisite for potent anti-MPER bnAb-mediated neutralization. However, in accordance with the latter proposed model, 10E8d was less potent than 10E8 even after increasing the local concentration via FcγRI, a finding that implies that lipid-binding may have a role beyond increasing the local concentration. However, it is possible that the extent of increasing the extent to which binding to FcγRI increased local antibody concentrations was not adequate, or that the slightly decreased affinity of 10E8d to MPER compared to 10E8 (Table 1) may have affected its neutralization potency on FcγRI-expressing cells.

**Fig. 5.**
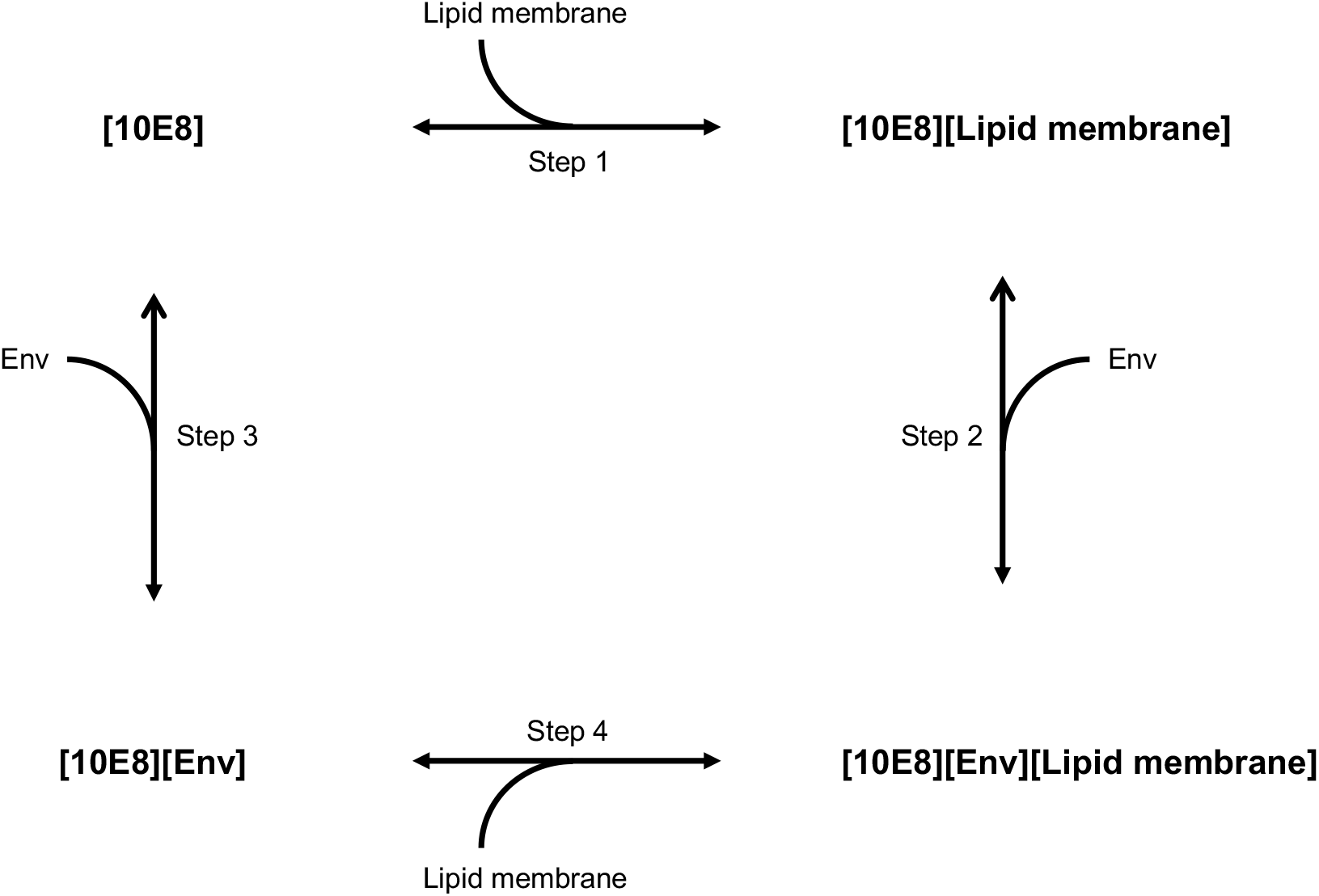
Schematic representation showing the relationship between 10E8, Env and lipid membrane on neutralization. Binding of 10E8 to lipid membrane (step 1) increases the local antibody concentration, enhancing neutralization by 10E8 upon binding to Env (step 2). Alternatively, 10E8 can bind first to Env (step 3) followed by binding to the lipid membrane (step 4), whereby lipid membrane-binding enhances neutralization by 10E8.

The MPER is a highly conserved motif within the HIV-1 Env gp41 subunit with high sequence conservation, so numerous attempts have been made to develop MPER-based vaccines (44). MPER-based fusion proteins as well as peptide or chimeric viruses have been developed and used to immunize animals, but these have elicited only non-neutralizing antibodies or antibodies with low potency and limited neutralizing breadth (45–50). Previous work has suggested that improved immunogenicity derived from displaying MPER on membrane lipids is based on the ability to promote a native-like conformation of the MPER (14, 44, 51). However, the exact conformation of MPER in this context is unknown. Additionally, autoreactivity of anti-MPER bnAbs as a precursor to immunologic host tolerance has been considered a hurdle to elicit potent anti-MPER bnAbs through immunization (52–54). Previous work has found that MPER-coupled liposomes elicited MPER-specific antibodies that recognize 10E8 epitopes but do not bind lipids, highlighting the fact that MPER specificity is not defined solely by lipid reactivity (55). Therefore, while lipids may be required to promote a native MPER conformation within an immunogen, lipid-binding may not be a requirement for bnAbs targeting MPER, where host tolerance is considered a hurdle to successful immunization.

Our work represents a new strategy to maintain potent neutralization by MPER-directed antibodies with decreased lipid-binding activity. Indeed, similar methods to improve neutralization potency by increasing local antibody concentrations have been validated. For instance, a 10E8v4/iMab targeting CD4 and MPER is in phase 1 clinical trials (NCT03875209), and it was proposed that the enhanced potency of this bispecific antibody relies on concentrating it at the site of viral entry via binding to CD4 (56, 57).

FcγRI is primarily found on monocytes and macrophages and anti-MPER antibodies could be protective for FcγRI-expressing macrophages and dendritic cells which are implicated to be among the first infection-susceptible cells that the virus encounters after exposure and early establishment of HIV-1 infection (58, 59, 68, 60–67). As well, HIV-1-infected macrophages rapidly spread the virus to autologous CD4^+^ T cells, at a proposed rate of one cell every 6 hours (63) and macrophages serve as a long-lived virus reservoir in chronic infection, posing a major hurdle to eradicating the virus in infected patients (69–73). On top of that, HIV-1 infection has been reported to prolong the life span of macrophages and to increase the mesenchymal migration of macrophages that leads to exposure of the virus to nearly all tissues including lung, liver, brain, urethra, lymph nodes, semen, gut, gut-associated lymphoid tissues and immune-privileged central nervous system (60, 74–82). Protecting these susceptible cells early during infection might limit the ability of the virus to take up long-term residence in niches that are difficult to reach with current therapeutics.

Infection prevention strategies using antibody or vaccine cocktails need to consider the sequence diversity of Env proteins as well as neutralization provided by antibody binding to Fc receptor. Currently, numerous clinical studies are being conducted to test antibody cocktails (83). For instance, several studies have demonstrated that the combination of bnAbs targeting different regions of Env protein showed additive neutralization *in vitro* (28, 31, 32, 84, 85). Candidate antibody combinations have also been described using mathematical modeling approaches (32). These findings highlight the additive effect of combining four bnAbs targeting different epitopes, compared to two- or three-antibody combinations. Antibody-mediated prevention (AMP) trials with passive transfer of the anti-gp120 CD4 binding site (CD4bs) bnAb, VRC01, have shown that the antibody failed to prevent viral rebound after antiretroviral therapy had been stopped (86, 87), though certain HIV-1 isolates were highly sensitive to VRC01 and could be neutralized during treatment. Finally, a non-human primate challenge study showed that the combination of two bnAbs, that alone had no effect, fully protected macaques against a mixed simian-human immunodeficiency virus (SHIV) challenge (88).

With a high level of interest in antibody combinations, it is important to understand how the various interactions synergize with each other when designing such strategies. In this work, the binding of 10E8 (via its Fc region) to FcγRI decreased its ability to synergize with NHR-targeting agents D5_AR and enfuvirtide (Figs. 3–4), possibly due to decreased accessibility of the NHR in the FcγRI-10E8-MPER-bound complex (Fig. 6). Alternatively, binding of 10E8 to FcγRI in the presence of NHR-bound agents may alter the orientation or position of 10E8, negatively impacting binding to either MPER or viral membrane, or both. A thorough understanding of the sequence of antibody-binding events will be critical for designing strategies to neutralize HIV-1.

**Fig. 6.**
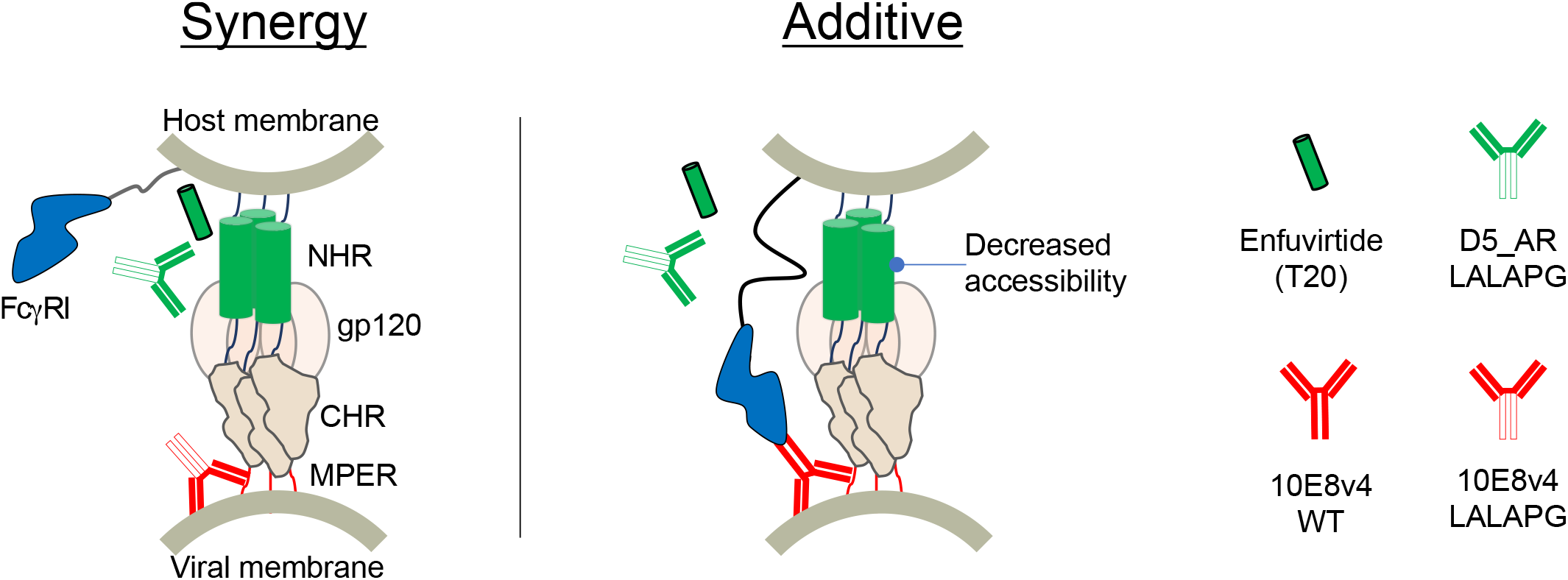
Hypothetical mechanism whereby steric hindrance affects synergy between NHR- and MPER-targeting agents. Synergistic interaction (left) between NHR-targeting agents (Enfuvirtide and D5_AR antibody) and MPER targeting antibody (10E8v4) turned additive when MPER-bound 10E8v4 also binds to FcγRI (right), possibly hindering accessibility of the NHR.

Overall, our findings suggest that lipid-binding activity is not a prerequisite for MPER-targeting bnAbs, motivate our efforts to design anti-MPER bnAbs whose local concentration can be increased without lipid-binding activity, and further contribute to our understanding of how to effectively target transiently exposed epitopes for the development of HIV-1 bnAbs and vaccines.

## Materials and Methods

### Antibody expression and purification

Genes encoding heavy and light chain of 10E8 and its variants (15, 19, 23) were synthesized by Integrated DNA Technologies (IDT). D5_AR IgG plasmids were obtained as described previously (34) and 10E8v4 (33) IgG plasmids were obtained from the NIH AIDS reagent program (APR-12866 and APR-12867 contributed by Dr. Peter Kwong). Sequences of heavy and light chain variable region of antibodies can be found in the Supplementary Information. PCR-based mutagenesis was used to introduce L234A/L235A/P329G mutations into the WT CH2 domain of IgG1 Fc to create LALAPG mutants. The gene encoding the LALAPG mutations were PCR-amplified using the following two primers: (HC_L234A_L235A-forward) 5’-CAGCACCTGAAGCCGCGGGGGGACCGTC-3’ and (HC_P329G_reverse) 5’-GATGGGGGCGCCGAGGGCTTTG-3’. The genes were then cloned into linearized pCMVR backbone using In-Fusion HD Cloning Kit Master Mix (Clontech). Antibodies were produced in Expi293F cells (Thermo Fisher Scientific) using FectoPRO (PolyPlus). Heavy and light chain plasmids were co-transfected at a 1:1 ratio. Cell cultures were incubated at 37°C under 8% CO_2_ with shaking at 120 rpm. The cells were harvested 4-6 days post-transfection by spinning at 5,000 × g for 10 min and filtering through a 0.22-μm filter. IgG were purified from supernatant using a 5 mL Mab Select Sure PRISM^™^ column (Cytiva) on the AKTA pure FPLC. The AKTA system was equilibrated with 1x phosphate-buffered saline (PBS, pH 7.4) and after binding of the antibodies followed by washing using PBS, the antibodies were eluted with 100 mM glycine (pH 2.8) into 1/10 volume of 1M Tris (pH 8.0). Eluates were buffer-exchanged into PBS and spin-concentrated using Amicon Ultra-15 100-kDa 15-mL spin concentrators (Millipore). The antibodies were filtered through a 0.22-μm filter and further purified using size-exclusion chromatography. The antibodies were purified on an AKTA FPLC using GE Superdex 200 increase 10/300 GL column (GE HealthCare) in 1× PBS.

### Fab generation

Fabs were generated from IgG via Lys-C-endopeptidase digestion following the manufacturer’s protocol (Wako). Briefly, 2 uL of Lys-C-endopeptidase was added to 500 uL of 2 mg/mL IgG in PBS with 1/10 volume of 1M Tris (pH 8.0) and digested for 1.5 hours at 37°C with moderate rotation. The digestion was stopped by adding 1/20 volume of 10% acetic acid. To ensure complete removal of undigested IgG and digested Fc, two-fold excess (6 mg of IgG/mL of resin) of protein A resin (Pierce) was added for 2 hours at room temperature with moderate rotation. Passthrough was collected and buffer-exchanged by spin concentration using Amicon Ultra-15 10-kDa 4-mL spin concentrators (Millipore).

### Synthesis of biotinylated MPER peptide

Biotinylated MPER peptide (KKKKWASLWNWFDITNWLWYIKLFIMIVGGKKK) was synthesized using standard Fmoc-based solid-phase peptide synthesis using CSBio instrument. NovaSyn TGR R resin (250 μmol, Novabiochem) was used and coupling was performed at 60°C for 15 min with 4-fold molar excess of amino acids. The peptide was biotinylated on the N terminus via coupling with biotin-polyethylene glycol (PEG)_4_-propionic acid (ChemPep). Dry peptide resin was cleaved using 94% trifluoroacetic acid, 2.5% water, 2.5% 1,2-ethanediol, and 1% triisopropylsilane at room temperature for 4 hours and precipitated in cold diethyl ether. Crude peptide was subsequently purified by reverse phase high-pressure liquid chromatography on a C_18_ semiprep column over an acetonitrile gradient with 0.1% trifluoroacetic acid and analyzed by liquid chromatography triple quadrupole mass spectrometry (Agilent).

### Biolayer interferometry

Biotinylated MPER peptide (30 nM) was loaded onto a streptavidin biosensor (Pall Fortebio) to a load threshold of 0.6 nm using an Octet RED96 system (Pall ForteBio). Ligand-loaded sensors were first baselined with octet buffer (0.5% [w/v] BSA in PBS containing 0.05% (v/v) Tween 20) followed by dipping into different concentrations (50 and 100nM) of 10E8 and its variants Fab for an association step (5 min) and returned to the baseline well for a dissociation step (10 min). Samples where biotinylated MPER peptide was loaded but did not associate with any 10E8 variants were used as a baseline subtraction. Response values (equilibrium dissociation constant (K_D_), K_on_ and K_off_) were calculated from Octet data analysis software. Response values obtained from different concentrations were averaged.

### Production of HIV-1 pseudotyped lentiviruses

HEK293T cells were transiently cotransfected with HIV-1 env plasmids (HXB2 and 25710) and psg3Δ Env backbone plasmids using a calcium phosphate transfection protocol, as described previously (34, 89–91). Briefly, 5 × 10^6^ HEK293T cells were seeded in 10-cm petri dish and incubated overnight at 37°C in a humidified atmosphere with 5% CO_2_. Backbone plasmid (20 μg) was mixed with Env plasmid (10 μg) and water for a final volume of 500 μL. Subsequently, 2× HEPES-buffered saline (pH 7, 500 μL, Alfa Aesar) was added dropwise to the mixture followed by 2.5 M CaCl_2_ (100 μL). The mixture was incubated for 20 min at room temperature and added dropwise onto the cells. 16 hours post-transfection, the medium was replaced with fresh medium. Supernatant was harvested 48 hours post-media replacement, centrifuged at 300 × g for 5 min and filtered through a 0.45-μm filter and stored at −80°C until further use. The backbone plasmid was obtained through the NIH AIDS Reagent program (catalogue number 11051) (92, 93).

### Cell culture

HEK293T and TZM-bl cells were maintained in Dulbecco’s modified eagle medium (DMEM) supplemented with 10% fetal bovine serum (FBS), 1% penicillin-streptomycin (Corning), and 1% L-glutamine (Corning) and incubated at 37°C in a humidified atmosphere with 5% CO_2_. TZM-bl cells were obtained through the NIH AIDS Reagent program from John C. Kappes and Xiaoyun Wu (92, 94–96). TZM-bl cells transduced to stably express FcγRI (TZM-bl/FcγRI) were maintained in DMEM supplemented with 10% FBS, 1% penicillin-streptomycin, 1% L-glutamine and blasticidin (Thermo Fisher Scientific) as described previously (25).

### Enzyme immunoassay

Nunc 96-well maxisorp plates (Thermo) were coated overnight at 4°C with 100 ng of FcγRI (Biolegend) in coating buffer (0.1M sodium bicarbonate, pH 8.6). Wells were blocked with 150 μL of ChonBlock blocking/sample dilution ELISA buffer (Chondrex) for 1 hour at 37°C. Anti-gp41 WT and LALAPG antibodies (0.0128~200nM) in ELISA buffer were added to the wells. After incubation for 2 hours at 37°C, plates were washed with 0.05% (v/v) tween 20 in PBS (PBST) three times followed by incubation with goat F(ab’)2 anti-human IgG (Fab’)2-horseradish peroxidase (Abcam) at a 1:10,000 dilution. The plates were washed again with PBST, followed by subsequent addition of Turbo 3,3′,5,5′ tetramethylbenzidine (TMB)-ELISA substrate solution (Thermo Fisher Scientific) and 2M sulfuric acid stop solution. Absorbance was measured at 450nm using a BioTek Synergy HT Microplate Reader.

### Flow cytometry

TZM-bl and TZM-bl/FcγRI cells were incubated with anti-gp41 WT and LALAPG antibodies (10 nM) in flow cytometry buffer (1% [w/v] BSA in PBS containing 0.05% [w/v] sodium azide) at 4°C for 1h and washed with flow cytometry buffer four times. Cells were then probed with Fluorescein (FITC)-Fab fragment goat anti-human IgG (1:50, Jackson ImmunoResearch) in flow cytometry buffer. After washing four times, the cells were sorted using BD AccuriTM C6 Plus (BD bioscience). 10,000 cells were detected per measurement and results were analyzed using Flowjo.

### Viral neutralization assay

TZM-bl and TZM-bl/FcγRI cells with tat-regulated luciferase reporter gene expression were used for quantification of viral infection and antibody neutralization as described previously (25, 26, 97). Briefly, 5 × 10^3^ TZM-bl or TZM-bl/FcγRI cells were seeded overnight in white-walled 96 well plates at 37°C in a humidified atmosphere with 5% CO_2_. The next day, the medium was aspirated without disturbing the cells and mixtures containing HIV-1 pseudotyped lentivirus, DEAE dextran (10 μg/mL) and anti-gp41 antibodies were added to the cells. After incubation for 48 hours, cells were lysed and luciferase activity was determined using BriteLite Plus reagent (Perkin Elmer). Relative luminescence unit (RLU) values were quantified using a Synergy HTX multimode reader (BioTek) and percent infection calculated as described previously (34).

### Analysis of synergy

Drug interactions for analysis of synergy were determined using the isobologram analyses. Isobologram analyses are based on dose-effect approaches which rely on mathematical framework known as Loewe additivity (98, 99). Loewe additivity builds on two concepts; dose equivalence principle (given an effect of dose a of drug A (d_a_), there is an equivalent dose b of drug B (d_b_) that gives the same effect, and reciprocally) and sham combination (d_b_ can be added to any other d_b_ to give the additive effect) and makes assumption that the drugs have a constant potency ratio (100). To construct isobologram for analysis of synergy, fixed-ratio dose-response neutralization curves for individual antibodies and antibody combinations were performed as described previously (101) with modification. Briefly, 100-fold ID_50_ values or antibody doses at 100% neutralization for individual antibodies were selected as starting concentrations. Antibodies were mixed by volume to volume ratio (1:0, 0:1, 1:1, 4:1 and 1:4) and serially diluted. The diluted antibody and antibody mixtures were tested for neutralization. From the dose-response neutralization curves, a dose-effect value of 50% neutralization was chosen to evaluate for antibody synergism. ID_50_ values for single antibodies in a given combination can be calculated using the equation:

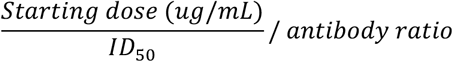

Isobologram was produced by plotting ID_50_ values for single antibodies in a given combination on an x-y coordinate graph. The straight line connecting the two ID_50_ values of the individual antibody is the locus of points that produce an additive effect (line of additivity).

## Acknowledgements

We thank Dr. Duo Xu, Dr. Benjamin Nikola Bell, Theodora Ulli Jordanka Bruun, and the Peter Kim Lab for fruitful discussion and useful comments on the manuscript. This work was supported by the National Institutes of Health under award number 5DP1AI15812502 (P.S.K), the Virginia & D.K. Ludwig Fund for Cancer Research (P.S.K.) and the Chan Zuckerberg Biohub (P.S.K.). S.K., and P.S.K. designed the project. S.K. performed the experiments. S.K., and M.V.F. generated reagents. S.K., M.V.F., and P.S.K. analyzed data and wrote the paper. All authors contributed to editing the manuscript.

## Supporting information captions

**Fig. S1.** Generation of 10E8 and its variants as Fab fragments

**Fig. S2.** Interaction of D5_AR and 10E8v4 antibodies with FcγRI

**Fig. S3.** D5_AR and 10E8v4 neutralize tier-1 and tier-2 HIV-1 pseudoviruses

**Table S1.** Neutralization potencies of D5_AR WT and 10E8v4 WT are potentiated by FcγRI

**Fig. S4.** FcγRI-bound 10E8d abolishes synergy with D5_AR

**Sequences.** Sequences of 10E8 variants and D5_AR

