## Supporting information for "Enhancing HIV-1 neutralization by increasing the local concentration of MPER-directed bnAbs"

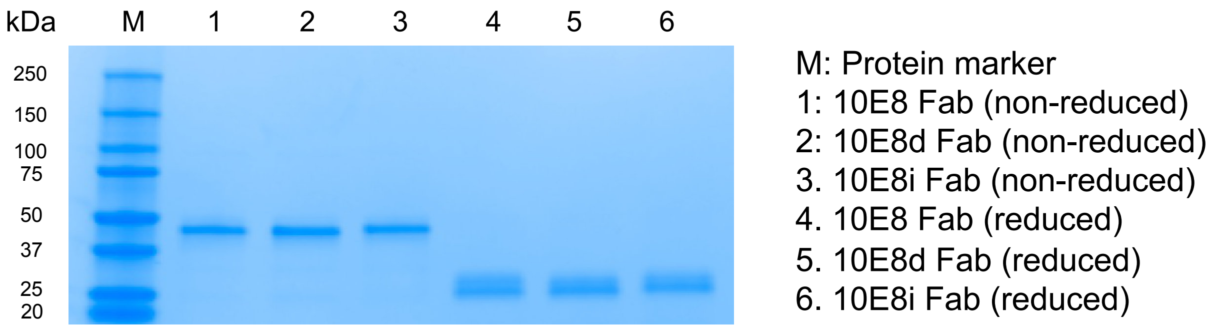


**Fig. S1.** Generation of 10E8 and its variants as Fab fragments. 10E8 and its variants (1 μg) were subjected to SDS-polyacrylamide gel electrophoresis and staining to visualize protein bands. Lane 1, non-reduced 10E8 Fab; lane 2, non-reduced 10E8d Fab; lane 3, non-reduced 10E8i Fab; lane 4, reduced 10E8 Fab; lane 5, reduced 10E8d Fab; lane 6, reduced 10E8i Fab. M, molecular weight marker.

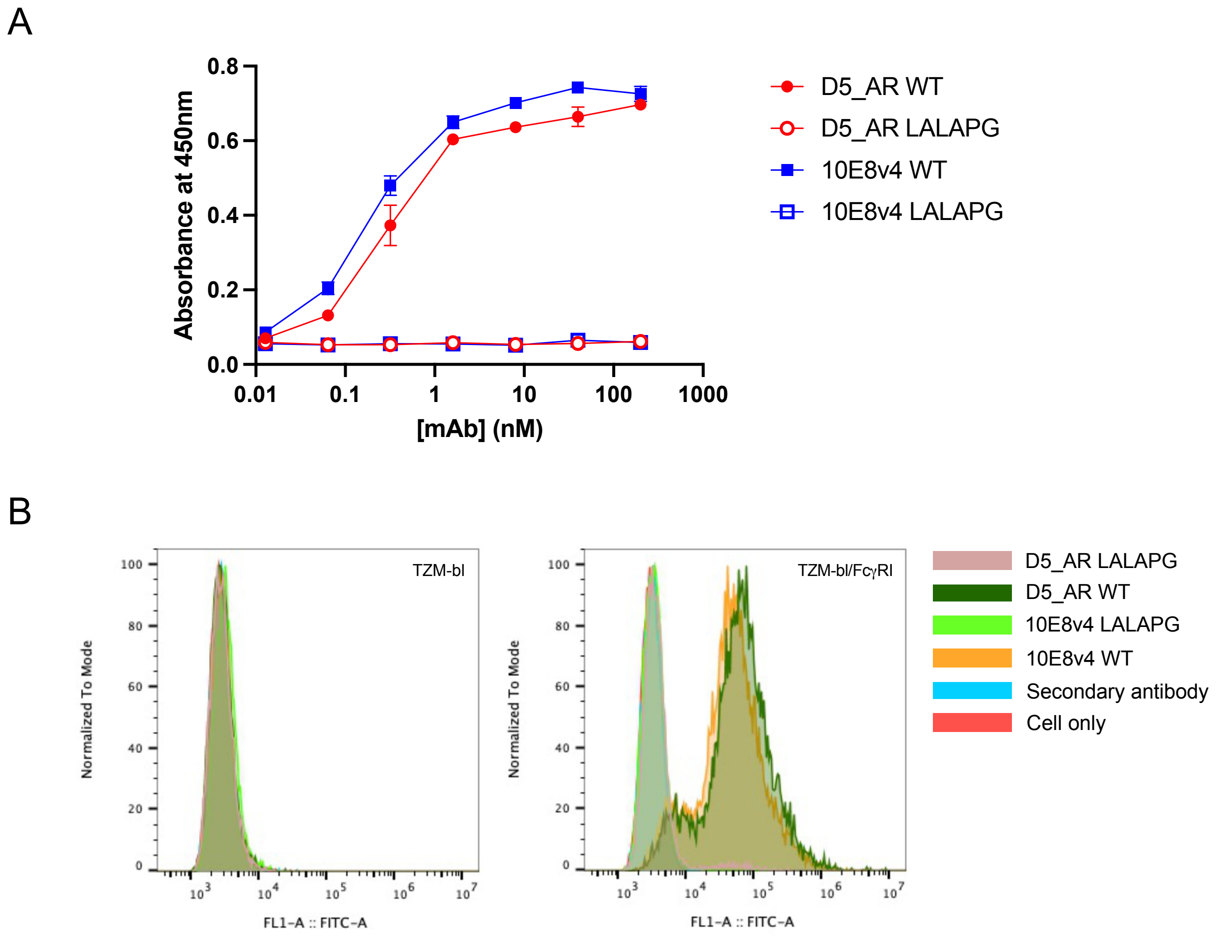


**Fig. S2.** Interaction of D5_AR and 10E8v4 antibodies with FcγRI. (A) Enzyme immunoassay of D5_AR and 10E8v4 WT and LALAPG mutants. FcγRI coated microtiter plates were incubated with various concentrations of D5_AR and 10E8v4 WT and LALAPG mutants (0.01-200 nM). Wells were subsequently probed with goat F(ab')2 anti-human IgG (Fab')2-HRP, and the amount of bound antibodies were determined using TMB substrate solution. Results are shown as mean ± standard deviations (SD) from triplicate experiments (technical replicates). (B) Flow cytometry of D5_AR and 10E8v4 WT and LALAPG mutants. TZM-bl cells (left) and TZM-bl cells expressing FcγRI (right) were treated with D5_AR and 10E8v4 WT and LALAPG mutants (10 nM) and probed with FITC Fab goat anti-human IgG (H+L).

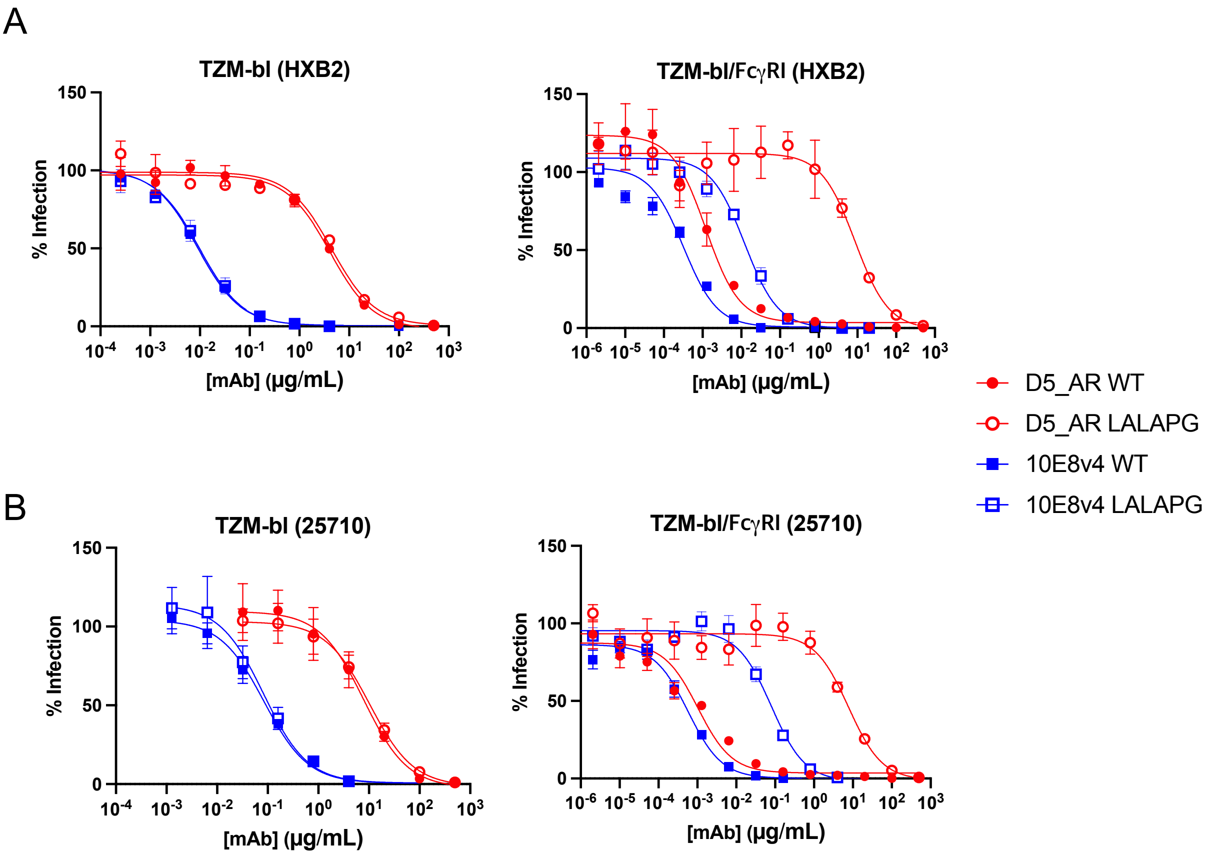


**Fig. S3.** D5_AR and 10E8v4 neutralize tier-1 and tier-2 HIV-1 pseudoviruses. (A) D5_AR and 10E8v4 WT and LALAPG antibodies inhibit infection of TZM-bl cells (left) and TZM-bl cells expressing FcγRI (right) against viruses pseudotyped with viral envelope glycoprotein (Env) from HXB2 (Tier 1). Results are shown as mean ± standard deviation (SD) from duplicate experiments (technical replicates). (B) D5_AR and 10E8v4 WT and LALAPG antibodies inhibit infection of TZM-bl cells (left) and TZM-bl cells expressing FcγRI (right) against viruses pseudotyped with Env from 25710 (Tier 2). Results are shown as mean ± SD from duplicate experiments.

**Table S1.** Neutralization potencies of D5_AR WT and 10E8v4 WT are potentiated by FcγRI

| Virus | Tier | Clade | Antibody | $ID_{50}$ (μg/mL) in TZM-bl | $ID_{50}$ (μg/mL) in TZM-bl/FcγRI | Fold enhancement by FcγRI |
|---|---|---|---|---|---|---|
| HXB2 | Tier 1 | Clade B | D5_AR WT | 4.1 | 0.0012 | 3400 |
| | | | D5_AR LALAPG | 4.4 | 8.7 | 0.51 |
| | | | 10E8v4 WT | 0.0089 | 0.00032 | 28 |
| | | | 10E8v4 LALAPG | 0.0097 | 0.012 | 0.81 |
| 25710 | Tier 2 | Clade C | D5_AR WT | 7.9 | 0.0010 | 7900 |
| | | | D5_AR LALAPG | 10 | 7.7 | 1.3 |
| | | | 10E8v4 WT | 0.084 | 0.00058 | 140 |
| | | | 10E8v4 LALAPG | 0.083 | 0.079 | 1.1 |

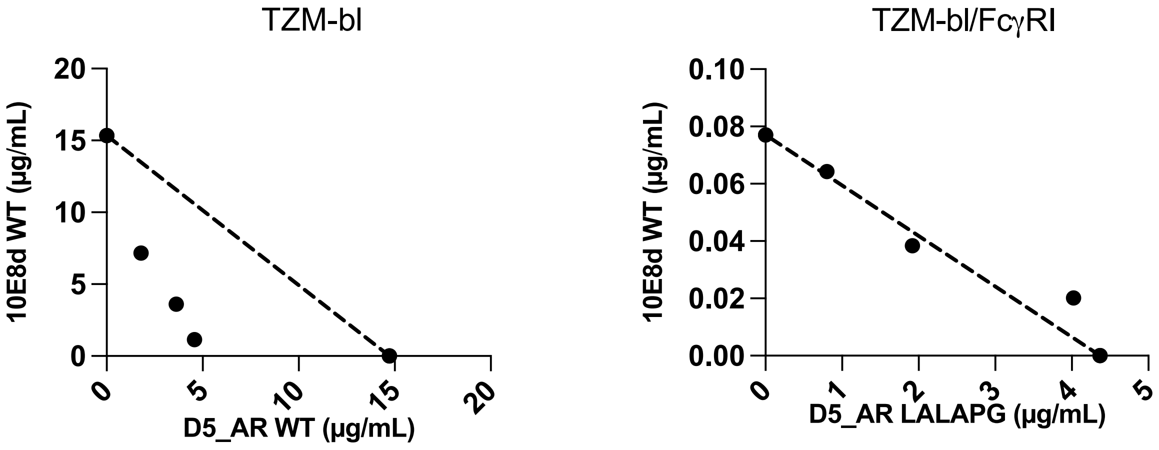


**Fig. S4.** FcγRI-bound 10E8d abolishes synergy with D5_AR. Isobologram analyses on the combination of 10E8d and D5_AR against viruses pseudotyped with Env from HIV-1 strains HXB2 in TZM-bl cells (left) and TZM-bl/Fc*y*RI cells (right). The dotted lines indicate lines of additivity. Results are shown as mean $ID_{50}$ performed in duplicates. Similar results were obtained in an independent repeat experiment. Data points below, along and above the line of additivity indicate synergy, additivity and antagonism, respectively.

**Sequences:**

**10E8 light chain**

SYELTQETGVSVALGRTVTITCRGDSLRSHYASWYQKKPGQAPILLFYGKNNRPSGVPDRFSGSASGNRASLTISGAQAEDDAEYYCSSRDKSGSRLSVFGGGTKLTVL

**10E8i light chain (mutations differing from 10E8 light chain underlined)**

SYELTQETGVSVALGRTVTITCRGDSLR**R**HYASWYQKKPGQAPILLFYGK**R**NRPSGVPDRFSGSA**R**GNRASLTISGAQAEDDAEYYCSSRDKSGSRLSVFGGGTKLTVL

**10E8d light chain (mutations differing from 10E8 light chain underlined)**

SYELTQETGVSVALGRTVTITCRGDSL**A**SH**A**ASWYQKKPGQAPILLFYGKNNRPSGVPDRFSGSASGNRASLTISGAQAEDDAEYYCSSRDKSGSRLSVFGGGTKLTVL

**10E8v4 light chain**

SELTQDPAVSVALKQTVTITCRGDSLRSHYASWYQKKPGQAPVLLFYGKNNRPSGIPDRFSGSASGNRASLTITGAQAEDEADYYCSSRDKSGSRLSVFGGGTKLTVL

**D5_AR light chain**

DIQMTQSPSTLSASIGDRVTITCRASEGIYHWLAWYQQKPGKAPKLLIYKASSLASGAPSRFSGSGSGTDFTLTISSLQPDDFATYYCQQYSNYPLTFGGGTKLEIK

**10E8 heavy chain**

EVQLVESGGGLVKPGGSLRLSCSASGFDFDNAWMTWVRQPPGKGLEWVGRITGPGEGWSVDYAAPVEGRFTISRLNSINFLYLEMNNLRMEDSGLYFCARTGKYYDFWSGYPPGEEYFQDWGRGTLVTVSS

**10E8v4 heavy chain**

EVRLVESGGGLVKPGGSLRLSCSASGFDFDNAWMTWVRQPPGKGLEWVGRITGPGEGWSVDYAESVKGRFTISRDNTKNTLYLEMNNVRTEDTGYYFCARTGKYYDFWSGYPPGEEYFQDWGQGTLVIVSS

**D5_AR heavy chain**

QVQLVQSGAEVRKPGASVKVSCKASGDTFSSYAISWVRQAPGQGLEWMGSIIPLFGT
AAYAQKFQGRVTITADESTSTAYMELSSLRSEDTAIYYCARDNPTFGAADSWGKGT
LVTVSS
